# Rapid range shifts in African *Anopheles* mosquitoes over the last century

**DOI:** 10.1101/673913

**Authors:** Colin J. Carlson, Ellen Bannon, Emily Mendenhall, Timothy Newfield, Shweta Bansal

## Abstract

Facing a warming climate, many tropical species-including the arthropod vectors of several infectious diseases-will be displaced to higher latitudes and elevations. These shifts are frequently forecasted for the future, but rarely documented in the present day. Here, we use one of the most comprehensive datasets ever compiled by medical entomologists to track the observed range limits of African malaria mosquito vectors (*Anopheles* spp.) from 1898 to 2016. Using a simple regression approach, we estimate that these species’ ranges gained an average of 6.5 meters of elevation per year, and the southern limits of their ranges moved polewards 4.7 kilometers per year. These shifts are consistent with the local velocity of climate change, and might help explain the incursion of malaria transmission into new areas over the past few decades. Confirming that climate change underlies these shifts, and applying similar methods to other disease vectors, are important directions for future research.

## Introduction

In the coming century, most scientific research projects a massive redistribution of global biodiversity. Today, the world is already +1.2° C warmer than in the pre-industrial period, and this transition is already underway: tropical species are spreading towards the poles, and species everywhere are tracking their thermal niche along elevational gradients. One foundational meta-analysis estimated that, to date, terrestrial species have been moving uphill at a pace of 1.1 meters per year, and to higher latitudes at a pace of 1.7 kilometers per year [1].

Among the millions of species on the move are some of the most consequential pathogens, disease vectors, and wildlife reservoirs that affect human health and economic development. For example, one study estimated that crop pathogens and agricultural pests were undergoing latitudinal shifts of 3 kilometers per year [2]. Similarly, the North American vector of Lyme disease, the deer tick *Ixodes scapularis*, has spread over 40 kilometers per year in the northeast [3,4]; the northern and elevational range limits of *Ix. ricinus* have expanded similarly rapidly in Europe [5–7]. In recent years, mosquito-borne diseases like malaria, dengue, and Zika virus have also expanded to new latitudes and elevations [8–10], and will continue to do so in the future, following the thermal limits on transmission set by their ectothermic vectors [11–13]. Some of these expansions have been facilitated by parallel global invasions of *Aedes aegypti* and *Ae. albopictus*, which have spread an estimated 250 and 150 kilometers per year, respectively; climate change will allow their spread to continue over the coming century, albeit at a slower pace [14,15].

However, surprisingly little is known about the impacts of climate change on the anopheline vectors of malaria, lymphatic filariasis, and O’nyong’nyong virus. Already, warming temperatures could have plausibly permitted expansions into highland east Africa [16]; some *Anopheles* species have become newly established in high-elevation sites in Latin America [17]; and a groundbreaking study recently found that in the Sahel, these mosquitoes can migrate hundreds of kilometers overnight, transported by wind currents [18]; but no systematic evidence exists that confirms range shifts are already underway in these species. Here, we track the geographic distributions of the primary malaria vectors (*Anopheles* spp.) in sub-Saharan Africa, and test the idea that over the last century, these species have moved southward (away from the equator) and upward (gaining elevation), consistent with hypothesized climate impacts.

## Methods

We revisit a recently published compendium of occurrence data for 22 species of *Anopheles* mosquitoes vectors of malaria in Africa [19]. While these data include a mix of finer taxonomy, we used the broadest possible definitions, treating *Anopheles funestus sensu lato* and *sensu stricto* as one species, and all members of the *Anopheles gambiae* complex - including *An. gambiae s.l., s.s.*, M form, and S form – as another single species.

In total, the dataset comprises over a century (1898 to 2016) worth of long-term, systematic entomological surveys from malaria programs, as well as other opportunistic data collected by researchers, gathered from a mix of peer-reviewed publications, technical reports, theses, and archival records. Records span more than one year at the majority of sampling sites (61%), covering an average of 8.5 years between the first and last presence record (Figure 1B). Parsed into unique spatiotemporal records, the dataset includes a total of 504,314 year-locality pairs, with an average of 22,923 records per species; while sampling fluctuates over time and increases during the Global Mariation Eradication Programme (1955-1969), the dataset spans the entire century with an incredible level of detail (Figure S1).

**Figure 1.**
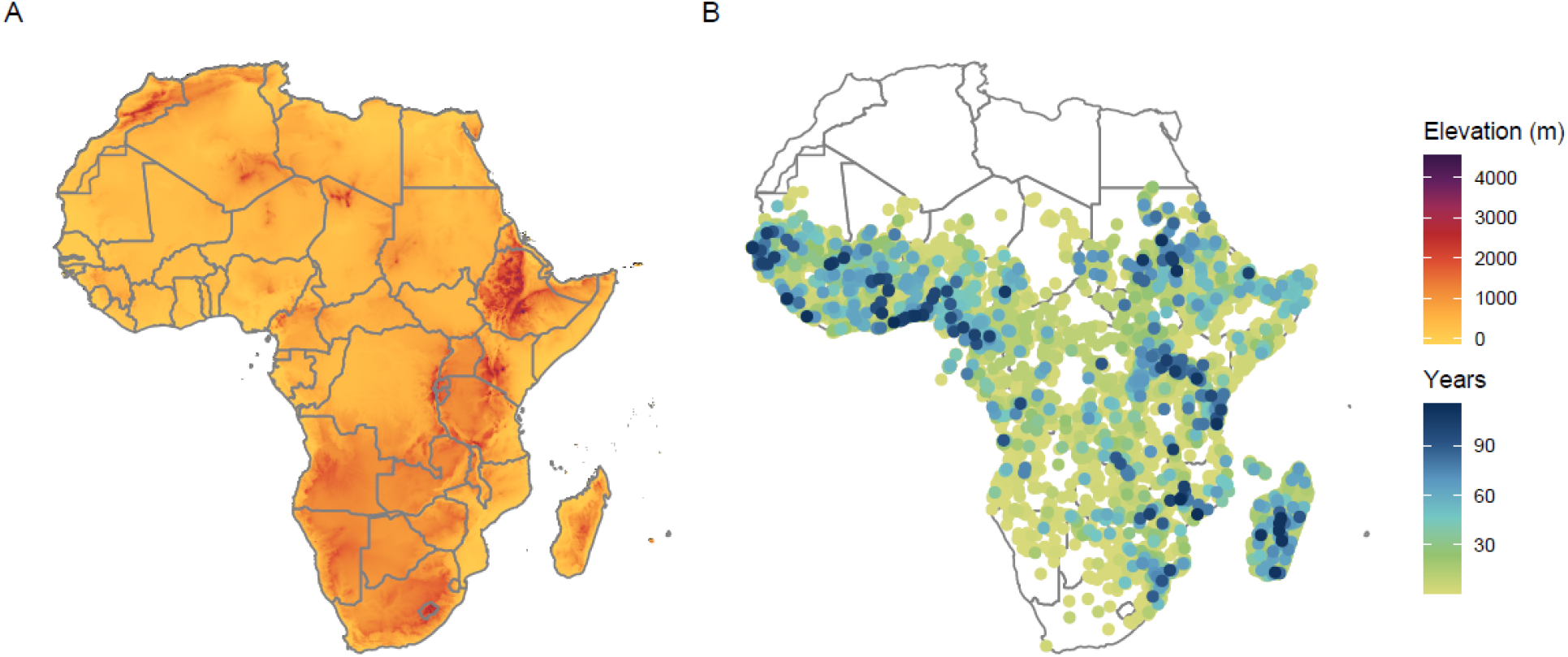
The elevational gradient in Africa (*left*); the sites of occurrence data, where color represents the maximum temporal span of observations (*right*).

For elevational data, we used the GTOPO30 global digital elevation model (DEM) downloaded as a 30 arc-second resolution grid for Africa from Data Basin (www.databasin.org; Figure 1A). We extracted elevation for each distinct occurrence record, using the ‘raster’ package in R version 3.3.2. In each year, we extracted the highest-elevation and southernmost records by species, limiting southernmost points to only those in the southern Hemisphere (to prevent any early years with incomplete sampling limited to west Africa from being included and inflating estimates).

For each of the 22 species of *Anopheles*, we used a simple linear regression with the ‘stats’ package to estimate change in elevational and latitudinal limits. We limited this analysis to cases with at least five or more unique values over time: with this cutoff, we were able to estimate latitudinal trends for 18 species, and elevational trends for 20 species. Latitudinal shifts were converted into approximate kilometers by assuming ~111km per degree of latitude.

## Results

In both elevational and latitudinal limits, we found a clear and unambiguous signal of long-term range expansion (Figure 2). We found that species’ southern range limits shifted at a pace of 0.042 decimal degrees (4.7 kilometers) each year, with 16 of 18 species exhibiting a significant trend (cutoff of p < 0.05). Elevational limits also shifted rapidly, with an average trend of 6.5 meters of altitudinal gain per year, and 18 of 20 species exhibiting a significant trend. All estimated elevational trends and most latitudinal trends (15 of 18) were positive, i.e., were consistent with the direction expected from climate-linked geographic range shifts. Finally, we found that the correlation between the two (*r* = −0.30) was insignificant (p = 0.25), suggesting that landscape-level patterns had a stronger influence on the pace of range shifts than variation among species’ intrinsic capacity for dispersal.

**Figure 2.**
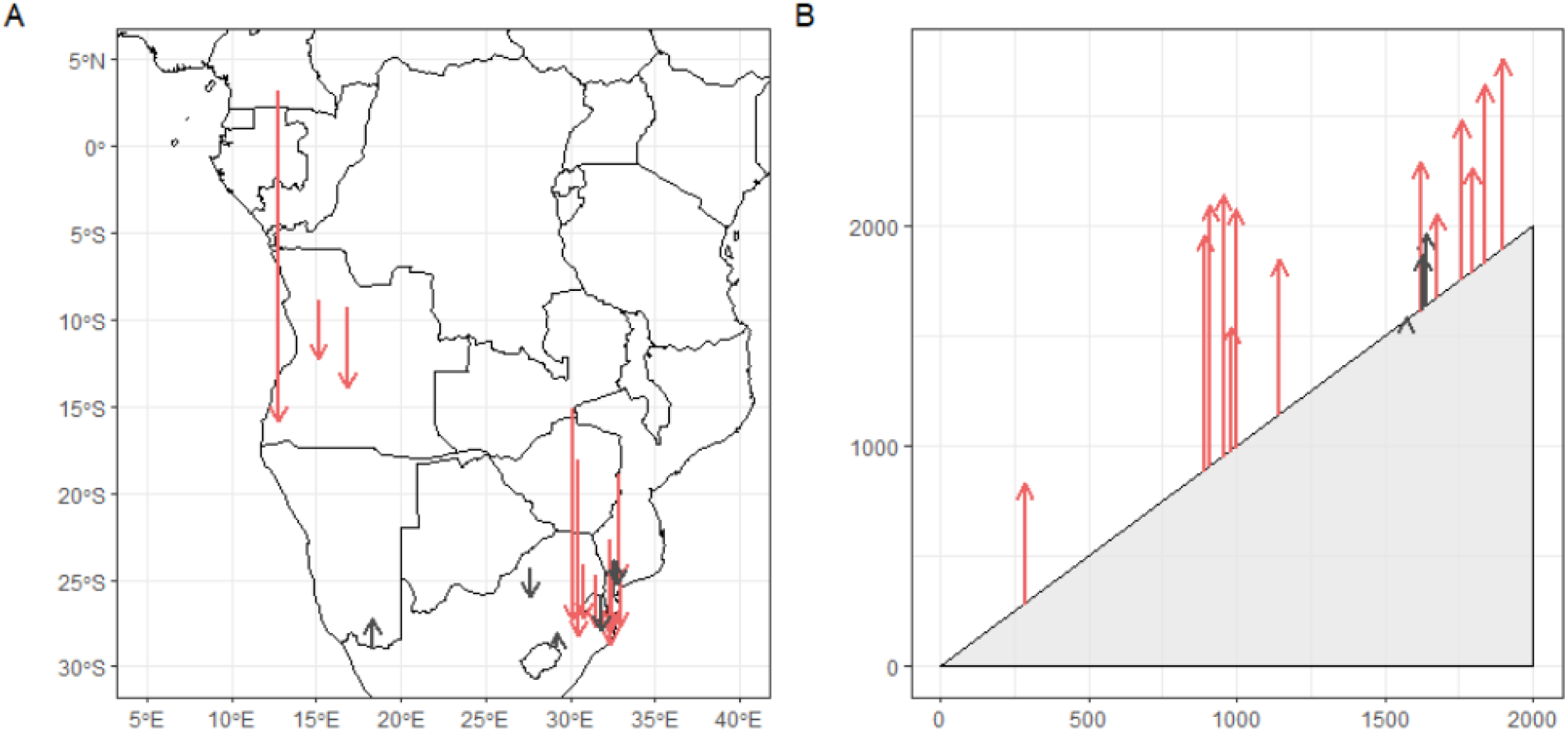
Estimated shifts in *Anopheles* species’ latitudinal and elevational maxima over the 20^th^ Century. *Left:* species’ southern maxima, where starting points are given at the longitude of the southmost point in the first half of the century (1900 to 1950), and the arrow shows the estimated latitudinal shift from 1900 to 2000 (chosen as a standardized unit for visualization, rather than the entire observation period, given that some species are sampled over slightly different intervals). *Right:* Elevational gain estimated from a linear model, 1900 to 2000 (y-axis), on a 1:1 elevational “gradient” (x-axis gives initial estimated elevational position). Red arrows indicate species for which temporal trends were statistically significant (*p* < 0.05).

## Discussion

We found clear evidence that *Anopheles* mosquitoes have undergone rapid range shifts over the 20^th^ century, challenging a long-standing assumption in historical epidemiology that mosquito ranges are mostly stationary over decades or centuries [20,21]. Our findings were consistent with expectations for the direction and pace of climate-linked range shifts, including previous estimates of climate velocity in sub-Saharan Africa [22]. Future work could build on these findings by using more sophisticated methods, such as spatiotemporal occupancy models [23], to formally test the explanatory power of climate change in these trends. If confirmed, the rapid expansion of *Anopheles* ranges—on average, over 500 kilometers southward and 700 meters uphill during the period of observation—would rank among the more consequential climate change impacts on African biodiversity that have been observed to date.

These findings could also suggest a new facet of the complex and contentious relationship between climate change and shifting malaria endemicity in Africa. The thermal limits of the *Plasmodium* parasite are well established [24,25], and readily superimposed onto climate projections; this mechanistic approach has suggested that malaria will spread into highland east Africa and expand at its southern range limits, but transmission will likely decrease as west and central Africa become prohibitively warm [13,26]. Beginning in the early 2000s, several studies have proposed that these impacts might already be observable in east Africa [27–29]. Others have disputed these conclusions, suggesting that they are irreconcilable with long-term progress towards malaria elimination, that trends in the region are better explained by lapsed control programs and growing drug resistance [30–33], and that climate change is inconsistent with long-term trends at the continental scale [34]. These debates—which remain unresolved—have focused nearly entirely on *P. falciparum* prevalence or incidence, and have rarely considered direct impacts of climate change on the mosquito vectors of the parasite.

If climate change has allowed *Anopheles* mosquitoes to invade once-protected colder areas, this might help explain observed changes in the altitudinal limits of malaria transmission [9], without presuming the veracity of a broader climate-driven, long-term increase in prevalence. Confirming this chain of causation would be an important step in resolving one of the longest-standing debates in climate and health research. More broadly, in the coming years, these sorts of direct links between climate, biodiversity change, and disease emergence will be increasingly important to quantify in real-time, not just to document a changing world but also to identify and address healthcare needs in newly-vulnerable populations.

## Data and Code Availability

No original data is used in this study. The study is fully reproducible with all code available on Github (github.com/cjcarlson/anophelev).

## Supplementary Information

**Figure S1.**
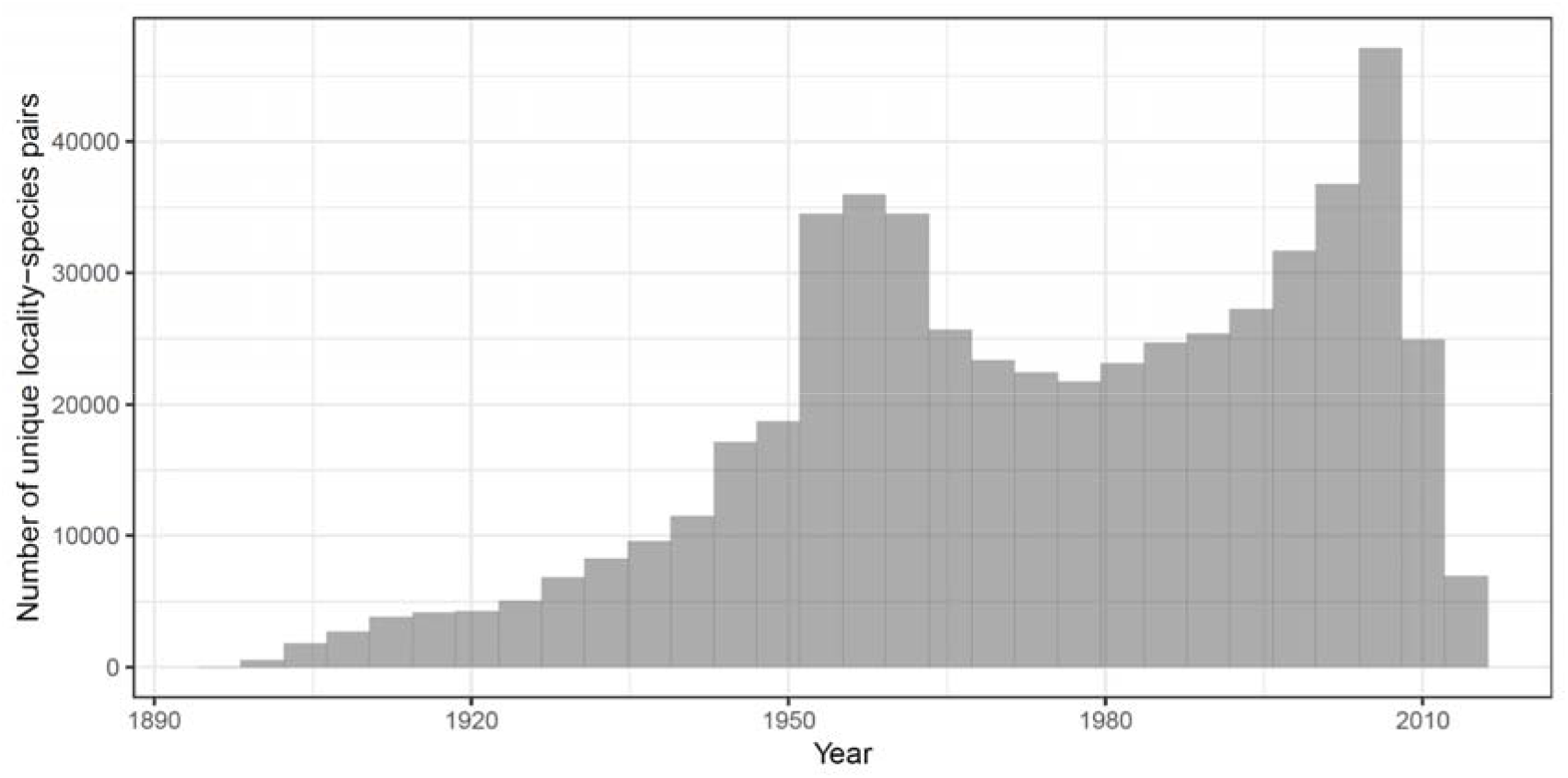
Sampling over time in the Kyalo *et al*. dataset.

